# Within-host population structure, migration, and parallel adaptive evolution of *Pseudomonas aeruginosa* in cystic fibrosis lung disease

**DOI:** 10.64898/2026.03.02.709079

**Authors:** David Ritz, Michelle E. Clay, Ted Kim, Rachel D. Van Gelder, Jayadevi H. Chandrashekhar, Alan J. Collins, Alix Ashare, Jay Goddard, Kenneth B. Hoehn, Daniel Schultz, Rachel J. Whitaker, Deborah A. Hogan

## Abstract

*Pseudomonas aeruginosa* infections in adults with cystic fibrosis (CF) are comprised of heterogeneous populations, most often tracing ancestry back to a single recent common ancestor. What is not clear is the physical spatial structure within the lung infection population, its stability over time and whether this physical structure leads to different evolutionary trajectories in different adaptive environments. To compare the *P. aeruginosa* populations across a single lung, we performed whole genome sequence analyses of 450 isolates recovered from lavage samples of the three different lobes of the right lung from a person with mild-to-moderate CF lung disease at three time points over the course of ∼1.5 years. We found that isolates fell into five distinct phylogenetic lineages with evidence for repeated translocation of isolates from different lineages across lobes and loss-of-function mutations in *lasR* and *mucA* were present in all 450 isolates. The well-resolved phylogenetic analyses revealed a structured population in which we find the coexistence of a slowly evolving lineage and more rapidly evolving lineages. There is also support for numerous migration events. Further, strong evidence for parallel adaptive mutations in multiple genes revealed distinct evolutionary paths affecting mucoid phenotypes and genetic variation in antibiotic resistance-associated pathways across coexisting populations within a single individual over time. These results provide an example of within-host evolution leading to microheterogeneity that may be useful to consider in future study of infection metapopulations dynamics over the course of chronic infection.

**IMPORTANCE:** Individuals with cystic fibrosis (CF) commonly have chronic lung infections that contain clonally derived *Pseudomonas aeruginosa* populations with genotypic and phenotypic diversity. This study describes a substantial dataset containing 450 isolates from different lobes of the right lung across three timepoints from an individual with mild-to-moderate CF lung disease. Some regional enrichment for specific lineages with parallel mutations among individual lobes of the lung was observed, but longitudinal analysis also demonstrated that compartmentalization is not strictly maintained and that isolates migrate between lobes of the lung over time. Perspectives on within lung evolution will be important for understanding the pathogen populations in chronic respiratory infections in CF and other diseases.

## INTRODUCTION

Pulmonary infection by *Pseudomonas aeruginosa* is known to correlate with increased lung disease severity and heightened mortality in individuals with cystic fibrosis (CF)^1–3^. Parallel selection for mutations in the same genes within and between CF infections supports the hypothesis that *P. aeruginosa* adapts to the CF lung environment over time^3^. For example, frequently observed loss-of-function mutations in the *lasR* gene, which encodes a quorum sensing regulatory transcription factor^4–7^ results in advantageous phenotypes including increased microoxic fitness^8^, improved growth^4^, and decreased elicitation of a host responses^9,10^. Another set of commonly mutated genes are those involved in the regulation of the production of alginate (e.g. *mucA* and *algU*), an exopolysaccharide that plays a protective role against a variety of environmental stressors^11–15^. Other common mutations include loss-of-function mutations in *mexZ* and other regulators involved in drug resistance, presumably because CF patients often receive frequent courses of antibiotics^16^.

Population scale studies of *P. aeruginosa* and other microbes associated with chronic infections have clearly established that phenotypically and genotypically heterogeneous populations persist in the CF lung over time^17–20^. Sequencing of *P. aeruginosa* isolates reveal that heterogeneity among strains often, but not always, converges to a single common ancestor, indicating emergence from a clonally derived population of *P. aeruginosa*^3,5,21–30^. Parallel independent evolution events have been observed in different lineages in a single patient^7^, suggesting the presence of a complex physically structured environment in the human lung. Lieberman et al. found similar preservation of diversity over time in *Burkholderia dolosa* and suggested that competing adaptive mutations might preserve diversity through clonal competition in addition to spatial structure^17^. An understanding of the spatial and temporal differences within single chronic infections will aid in devising suitable treatment regimes. Towards increasing this understanding, Jorth et al.^21^ used lung explants from CF subjects with severe disease to provide phenotypic and genotypic evidence for the physical separation of *P. aeruginosa* populations and their independent evolution. While this study clearly supports the model that physical structure limits migration between lobes, it is not clear if subpopulations are stably associated with specific regions over longer periods of time.

To contribute to the understanding of the evolutionary dynamics of *P. aeruginosa* over space and time, we examined *P. aeruginosa* populations derived from bronchoalveolar lavage (BAL) fluid. Four hundred and fifty isolates from BAL fluid collected from different lobes of the right lung from the same individual with mild CF lung disease at three time points were analyzed by whole genome sequencing. Visualization of the phylogenetic relationships among these isolates showed the presence of five major clusters. Members of each cluster were enriched in specific lobes at a single time point, but there was evidence for migration between lobes between time points followed by expansion of specific subpopulations. Further, there were differences in mutation rates in different regionally-localized subpopulations. While some genetic characteristics associated with CF, such as nonsense mutations in *lasR* and *mucA* genes, were observed in all isolates, there were many instances of parallel evolution events associated with alginate production and drug resistance. Together, these data reveal a dynamic, spatially structured population in which migration and parallel adaptation collectively shape within-host *P. aeruginosa* evolution over time.

## RESULTS

### Characterization of reference genome from a timepoint 1 isolate

*P. aeruginosa* isolates (n = 527) were obtained from BAL samples collected from the right upper, middle, and lower lobes (RUL, RML, and RLL) of an individual with mild-to-moderate CF lung disease at three time points: Timepoint 1, Timepoint 2 (151 days later), and Timepoint 3 (353 days after Timepoint 2) **(Fig. 1A)**. The subject was an adult male with CF, 30-35 years of age at the time of sampling, with an average lung function of 94% predicted forced expiratory volume (FEV1) across the study period. During pulmonary exacerbations, FEV1 declined to approximately 75%. Clinical sputum cultures documented chronic colonization by both mucoid and non-mucoid *P. aeruginosa* for more than eight years prior to the first BAL collection. Throughout the study period, the subject received multiple antimicrobial treatments, including azithromycin, ceftazidime, tobramycin, ciprofloxacin, vancomycin, piperacillin-tazobactam (Zosyn), doxycycline, and trimethoprim-sulfamethoxazole (Bactrim) as described in detail below.

**Figure 1.**
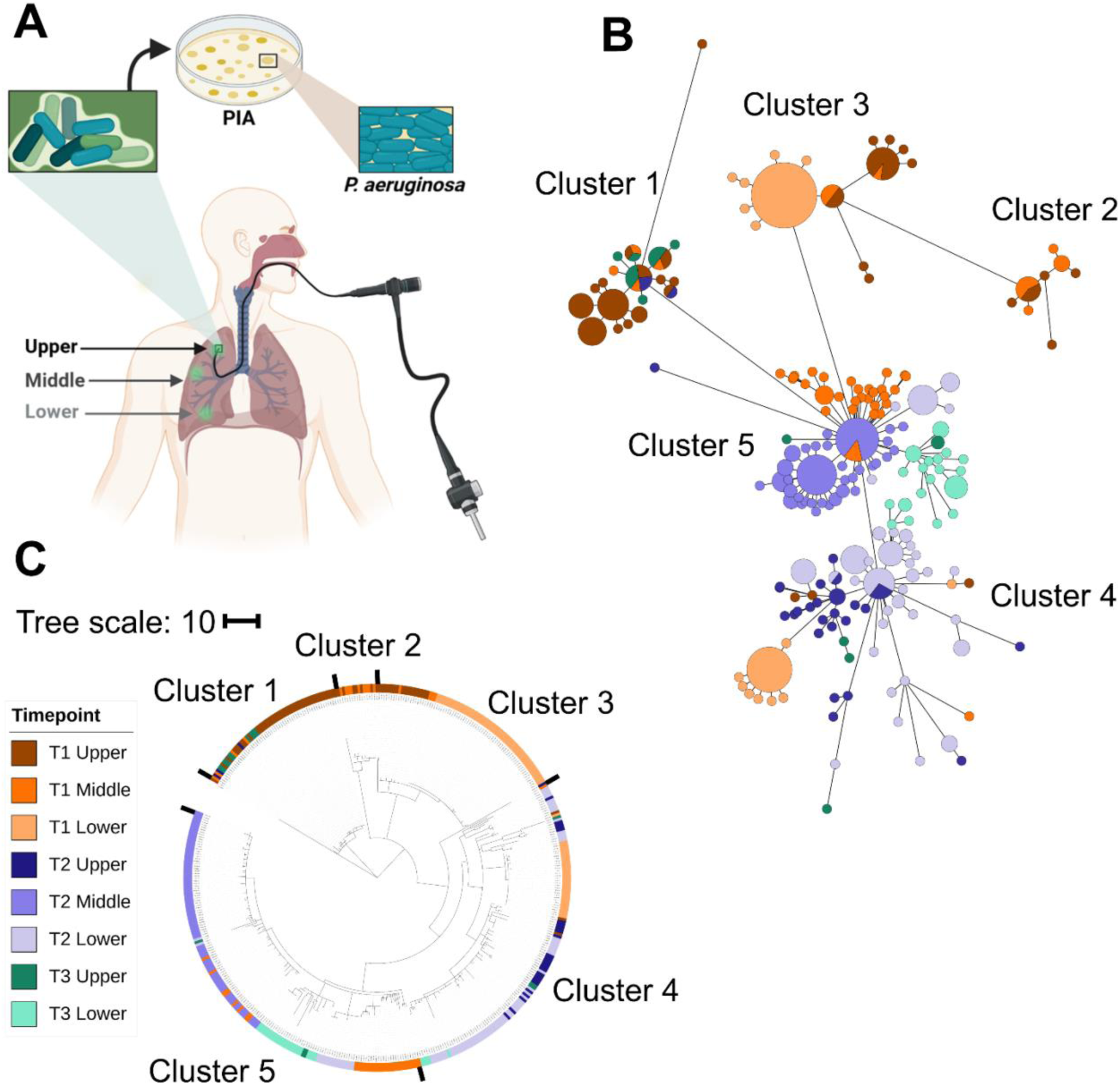
Diverse *P. aeruginosa* strains were isolated from the CF subject’s right lung. **(A)** The methodology for collection for bronchoalveolar lavage (BAL) fluid from the upper, middle, and lower lobes of the subject’s right lung at three timepoints and subsequent isolation of *P. aeruginosa*. **(B)** GrapeTree represents relationships between sets of identical haplotypes. The size of the grape circle represents the proportion of strains with that haplotype from each sample, and the edges connect haplotypes. The length of the edges represents the number of SNPs between nodes. The three timepoints are: Timepoint 1, Timepoint 2 (151 days later), and Timepoint 3 (353 days after Timepoint 2) **(C)** Rooted circular phylogenetic tree of 450 BAL isolates that includes the ST499 outgroup.The colored bars indicate the isolates’ origin and time points of the BAL. The branch length is equivalent to the number of detected SNPs.

To study the population structure and distribution of genotypes over time, the 527 strains were sequenced with short-read Illumina technology. To identify single nucleotide polymorphisms (SNPs) that were unfixed in the population, reads were aligned to an internal reference genome created by sequencing the DNA from isolate T1_LL_B5 using long-read Nanopore technology with polishing using Illumina short read sequences. All isolates belonged to multilocus sequence type (MLST) ST499, consistent with a single common infection origin. Isolate T1_LL_B5 harbored approximately 3,000 SNPs relative to a publicly available genome for another ST499 strain (ERR364087). The ST499 comparator and T1_LL_B5 sequences were more closely related to the epidemic strain LESB58 than to the commonly used laboratory strains PAO1 and PA14 **(Fig. S1)**.

### Detection of variable SNPs in the single-subject isolate collection

After read normalization and alignment of the 527 isolates to the T1_LL_B5 reference genome, variable positions were identified across isolates using breseq^31^. Further, both genomes and positions were then filtered to exclude low confidence data using read depth and consensus score metrics as described in detail in the methods. The 450 isolate genomes that remained after filtering included 45 to 201 isolates per timepoint and 121 to 196 isolates per lobe **(Table 1).** Across these strains, there were 663 high confidence variable SNPs **(Supplementary Data 1)** which included 401 non-synonymous SNPs, 198 synonymous SNPs, and 64 intergenic SNPs. In addition, insertions and deletions (indels) were recorded across the collection **(Supplementary Data 2)**.

**Table 1.** *P. aeruginosa* isolated from right lung of person with CF. . Data stratified by number of isolates per lung lobe used in the analyses.

| Timepoint | No. of isolates |  |  |
| --- | --- | --- | --- |
|  | Upper lobe | Middle lobe | Lower lobe |
| T1 | 73 | 52 | 85 |
| T2 | 32 | 81 | 82 |
| T3 | 16 | - | 29 |

We first assessed the phylogenetic relatedness of the population using an unrooted grape tree which facilitates the visualization of phylogenetic relationships in the context of metadata such as location and time point **(Fig. 1B)**. We observed the population resolved into five major “clusters”, with isolates from the three time points and the three sampled lung regions within each cluster. The clusters are also indicated on a phylogenetic tree shown in **Fig. S2**. In a rooted phylogenetic tree **(Fig. 1C)** that included the ST499 comparator as an outgroup, we observed that the isolates that predominated in the upper lobe within Cluster 1 were most closely related to the outgroup. We also observed enrichment of derived lineages within specific regions at each time point as well as strains from different coexisting within regions suggesting migration among lobes.

### Isolates have heterogeneous evolution rates and migration patterns

We used the phylogenetic analysis of isolate genomes to reveal dynamics of *P. aeruginosa* evolution and migration within the right lung. If *P. aeruginosa* was evolving over time, isolates at T3 would be more genetically diverged from the most recent common ancestor than isolates at T1 (**Fig. 1C**).To estimate the average rate of this evolution, we used a root-to-tip regression of genetic divergence against sample date (**Fig. 2A**)^32^. The slope of this regression line indicated an overall rate of ∼0.02 mutations per day across the study, significantly greater than zero when assessed using a permutation test (P < 1×10^-4^). This rate of evolution was not uniform, however. For example, isolates within Cluster 1 showed no increase in divergence, and therefore no evidence of evolution over the 485 days between T1 and T3 (**Fig. 2A**). These Cluster 1 strains were the least diverged from the most recent common ancestor and were predominantly localized to the upper lung lobe. These results indicate that *P. aeruginosa* measurably evolved during the study, but that this rate of evolution was heterogeneous among regions and strains within the right lung.

**Figure 2:**
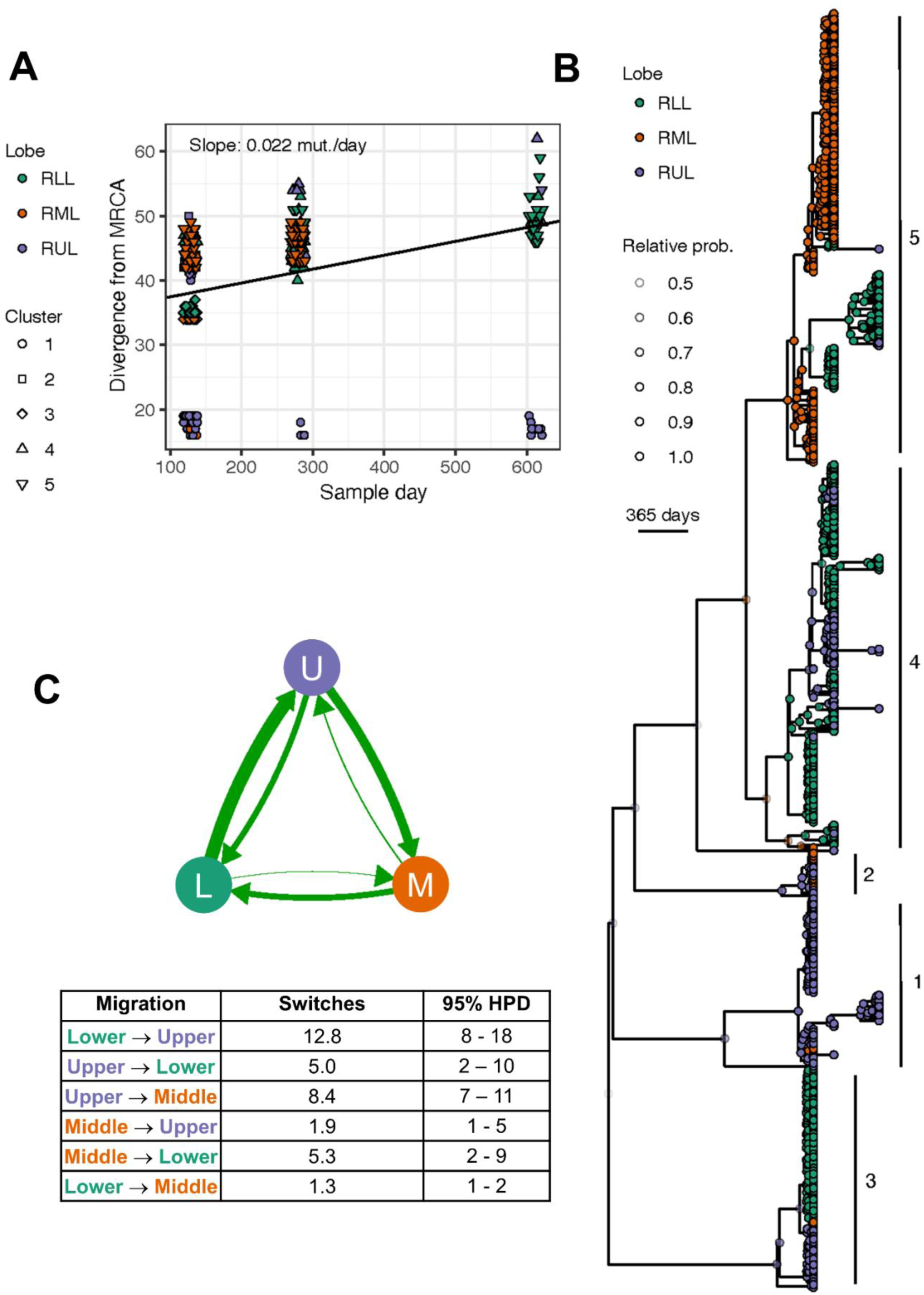
Migration and mutational rates depend on lung lobe. **(A)** Estimated number of migration events between lobes along time trees. Arrow widths are proportional to the mean number of switches (parent nodes with different lobes than child nodes) in each direction along trees sampled from the posterior distribution. The mean number of migrations and their corresponding 95% highest posterior density (HPD) interval are included as a table. **(B)** Time-resolved phylogenetic tree (maximum clade credibility tree) of the CF isolates. Internal nodes are labeled by their estimated location and shaded by its posterior probability. Tips are further labeled by cluster number. **(C)** Root-to-tip regression of genetic divergence of each isolate from the recent common ancestor (MRCA) compared to sample date. Horizontal jitter added to reduce overplotting. The slope of this regression line is the estimated average rate of evolution over the sample interval. The color code is not consistent between Fig. 2 and the other Figures.

To link the spatial and temporal evolution of *P. aeruginosa* within this subject’s right lung, we inferred time-resolved phylogenies from the CF isolate genomes **(Fig. 2B)**. To do so, we used an uncorrelated lognormal relaxed molecular clock model implemented in BEAST2^33,34^. We estimated the date of the most recent common ancestor of all sampled isolates as ∼4.1 years prior to T1 (95% HPD: 1.8 – 7.2 years), indicating years of chronic infection compatible with the initial *P. aeruginosa* infection beginning more than 8 years prior to the study. Major genetic clusters 1-5 were identifiable within the maximum clade credibility tree **(Fig. 2B)**. To infer patterns of migration among lobes, we modelled location as a discrete trait using a continuous time Markov model (see Methods)^33,35^. Across the trees sampled from the posterior, we observed a mean of 32.9 pairs of parent/child nodes where the child node had a different location from the parent, representing a migration event between lobes **(Fig. 2C)**. These switches most frequently occurred from the lower to upper lobe, followed by upper to middle, and middle to lower. Meanwhile, migration events in the opposite directions were much less frequent. These results suggest a cyclical pattern of migration in which *P. aeruginosa* lineages move from the upper lobe through the middle and lower lobes and subsequently return to the upper lobe.

Pairwise fixation index analysis (F_ST_), which combines both genetic distance and abundance in all pair wise comparisons between samples, also revealed dynamic spatial and temporal restructuring of the bacterial population across lung lobes (**Table S1**). For the most part samples are highly differentiated (F_ST_ >0.1) likely showing the clonal nature of local growth captured by BAL sampling. However, the few pairs of samples with very low differentiation (<0.07) supported the evidence for mixing between regions over time. At T1, moderate differentiation among upper, middle, and lower lobes (FST ≈ 0.35–0.37) indicated spatial compartmentalization, consistent with partially isolated intra-lung niches. At T2, this structure was dramatically altered: the upper and lower lobes became highly similar (FST = 0.069), suggesting substantial bacterial mixing or redistribution between these regions. In contrast, the middle lobe at T2 was strongly differentiated from both contemporaneous lobes (FST = 0.60–0.71). By T3, spatial structure re-emerged (T3 upper vs. lower FST = 0.477), and notably, the T3 upper lobe population was nearly identical to the T1 upper lobe population (FST = 0.058), suggesting persistence or re-expansion of the slow growing ancestral lineage following the transient T2 restructuring event. It is worth noting that intermediate F_ST_ values were observed between the middle lobe at T1 and both the lower and upper lobes at future time points suggesting there was some migration from middle lobe in both directions over time while the middle lobe is dominated by single type. Without a T3 sample for the middle lobe, the impact of mixing in this region was not assessed. Together, these data support a model in which intra-lung bacterial migration is episodic rather than constant, with temporary breakdown of spatial barriers.

### Predicted adaptive mutations within the population

The evolution of *P. aeruginosa* during the infection resulted in mutations in several mechanisms known to be involved in adaptation to the human host. Missense and nonsense mutations were found in 333 unique PAO1 ORFs, 91% of which were only mutated in single strains. Twenty-two genes had mutations in two or more lineages, indicating parallelism and potentially adaptive evolution **(Supplementary Data 1)**. In this section, we focus on specific pathoadaptive loci impacting phenotypic characteristics that have been well characterized and are medically relevant.

#### A lasR mutation was fixed in the population

An analysis of the colony morphologies of the isolates found that subsets of the population had a sheen due to the accumulation of 2-heptyl-4-hydroxyquinoline (HHQ), which is very common in strains with loss-of-function mutations in *lasR*^4^. While *lasR* was not identified as a gene with unfixed SNPs in the population, comparison of the *lasR* gene sequence in all the isolates to the *lasR* sequence from strain PAO1 found that all 450 isolate genomes contained a nonsense mutation at amino acid Gln45 due to a C135T nucleotide change in the *lasR* gene which led to premature termination. These data suggest that a *lasR* mutation was either present in the original infecting strain or had come to dominate the population in the right lung prior to our studies. As not all isolates had the *lasR* colony phenotype, we searched for mutations in other genes that could impact HHQ production. We found only one isolate with a high confidence missense mutation in *pqsE*, which regulates the production of HHQ and its derivative PQS, but no SNPs in other *pqs* genes **(Supplementary Data 1)**. Future studies would be required to determine how other loci mutated in this study affect HHQ accumulation and *lasR* phenotypes.

#### Evolution and regional differences of the mucoid phenotype

We observed that a subset of isolates exhibited a mucoid phenotype associated with increased production of the exopolysaccharide alginate. All 450 isolate genomes carried an identical mutation in *mucA* consisting of a single-nucleotide deletion after amino acid position 143 **(Fig. 3A)**. This frameshift (fs) altered residues 144–146 from Ala-Pro-Gly to Arg-Arg-Arg and introduced a premature stop codon (*mucA*144fs), resulting in truncation of the MucA protein. Loss-of-function mutations in the MucA anti-sigma factor are well known to activate alginate production due to increased AlgU activity^36^. Additional mutations were observed in other components of the alginate biosynthetic pathway. Sixteen isolates from the right upper and middle lobes at timepoint 1 formed a distinct group carrying mutations in both *algG* (R11H) and *algL* (S203Y) **(Fig. 3A)**, suggesting alternative genetic routes to modulate alginate production.

**Figure 3:**
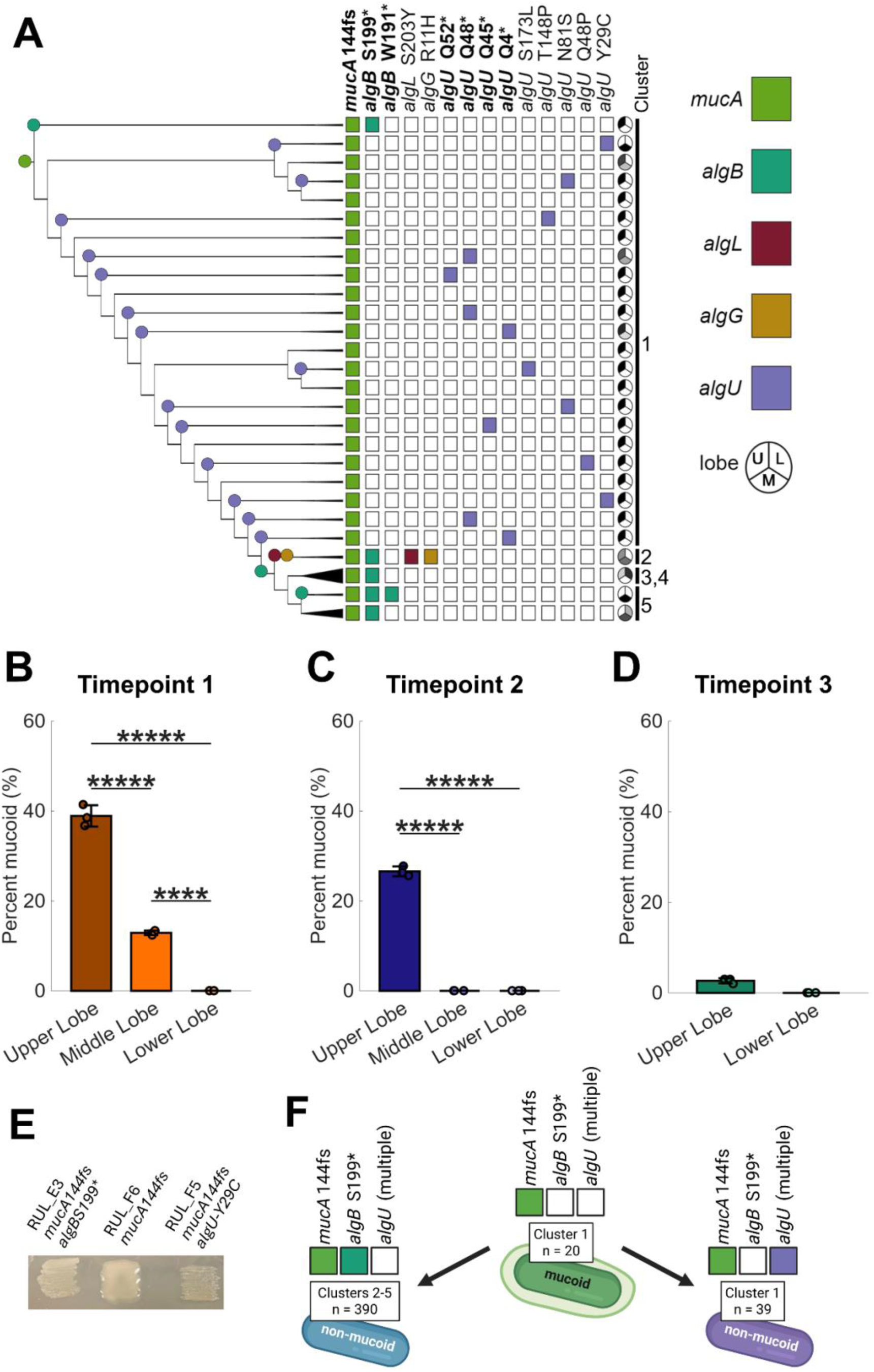
Adaptive mutations cause a variable mucoid phenotype. **(A)** Mutations affecting alginate production are marked in the phylogeny tree of the *P. aeruginosa* isolates. Adjacent strains with identical mutations were collapsed into the same leaf for clarity, with the size of the wedges representing the number of isolates collapsed into each leaf. Colored dots represent where each mutation likely occurred in the phylogeny tree. Cluster numbers are adjacent to the lobe pie graphs. Bold x-axis labels represent nonsense or frameshift mutations likely leading to loss of protein function. **(B-D)** The mucoid phenotype was quantified for strains isolated at **(B)** timepoint 1, **(C)** timepoint 2, and **(D)** timepoint 3, with comparisons between the isolate origins. Asterisks denote significance levels for pairwise comparisons between groups: ***** p < 10^-5^, **** p < 10^-4^, *** p < 10^-3^, ** p < 10^-2^, and * p < 0.05. **(E)** Representative mucoid patches of isolates with either a *mucA* frameshift and *algB* nonsense mutation, only a *mucA* frameshift mutation, or a *mucA* frameshift and *algU* mutation, respectively. **(F)** A frameshift mutation in *mucA* after amino acid 143 is fixed in the population and confers an alginate-producing phenotype in Cluster 1 strains. Subsequent mutations in *algU* or *algB* restore the non-alginate phenotype. One outlier strain from Clade 1, which carries the *algB* S199* mutation, is not represented in this schematic and corresponds to the first row in Fig. 3A.

Regional and temporal differences in mucoidy were evident across the isolate collection. Mucoid isolates were most prevalent in samples from the right upper lobe, and the overall frequency of mucoidy declined over the course of the study **(Fig. 3B–D)**. In mucoid strains with low MucA activity, subsequent mutations in *algU* can cause reversion from mucoid to the non-mucoid phenotype^37^. Consistent with this, our analysis of unfixed mutations identified nine non-synonymous mutations in *algU*, all confined to Cluster 1 isolates, including four nonsense mutations and five missense substitutions **(Fig. 3A)**. In contrast, *algU* remained functional in all other isolates. Notably, outside of Cluster 1, all isolates carried a nonsense mutation in *algB* (S199*), a positive regulator of *algD*, which is a GDP-mannose 6-dehydrogenase essential for alginate biosynthesis^38^. In addition, a small group of isolates in Cluster 5 encoded an additional nonsense mutation in AlgB (W191*), resulting in a second independent truncation event **(Fig. 3A)**. Phenotypic assessment of isolates harboring *algU* or *algB* mutations in the fixed *mucA* loss-of-function background shows loss of mucoidy **(Fig. 3E)**, indicating that a fixed *mucA* frameshift mutation initially predisposed the population to alginate production, while subsequent lineage-specific mutations in *algU* and *algB* drove repeated reversions to the non-mucoid phenotype **(Fig. 3F)**.

#### Efflux pump mutations are not predictive of antibiotic resistance phenotypes

Our analysis of unfixed mutations found a wealth of non-synonymous mutations in genes encoding multidrug efflux pumps implicated in antibiotic resistance. In many instances, the same genes acquired missense or nonsense mutations in multiple lineages from distinct clusters, suggesting parallel adaptation to antibiotic treatments **(Fig. 4A)**.

**Figure 4:**
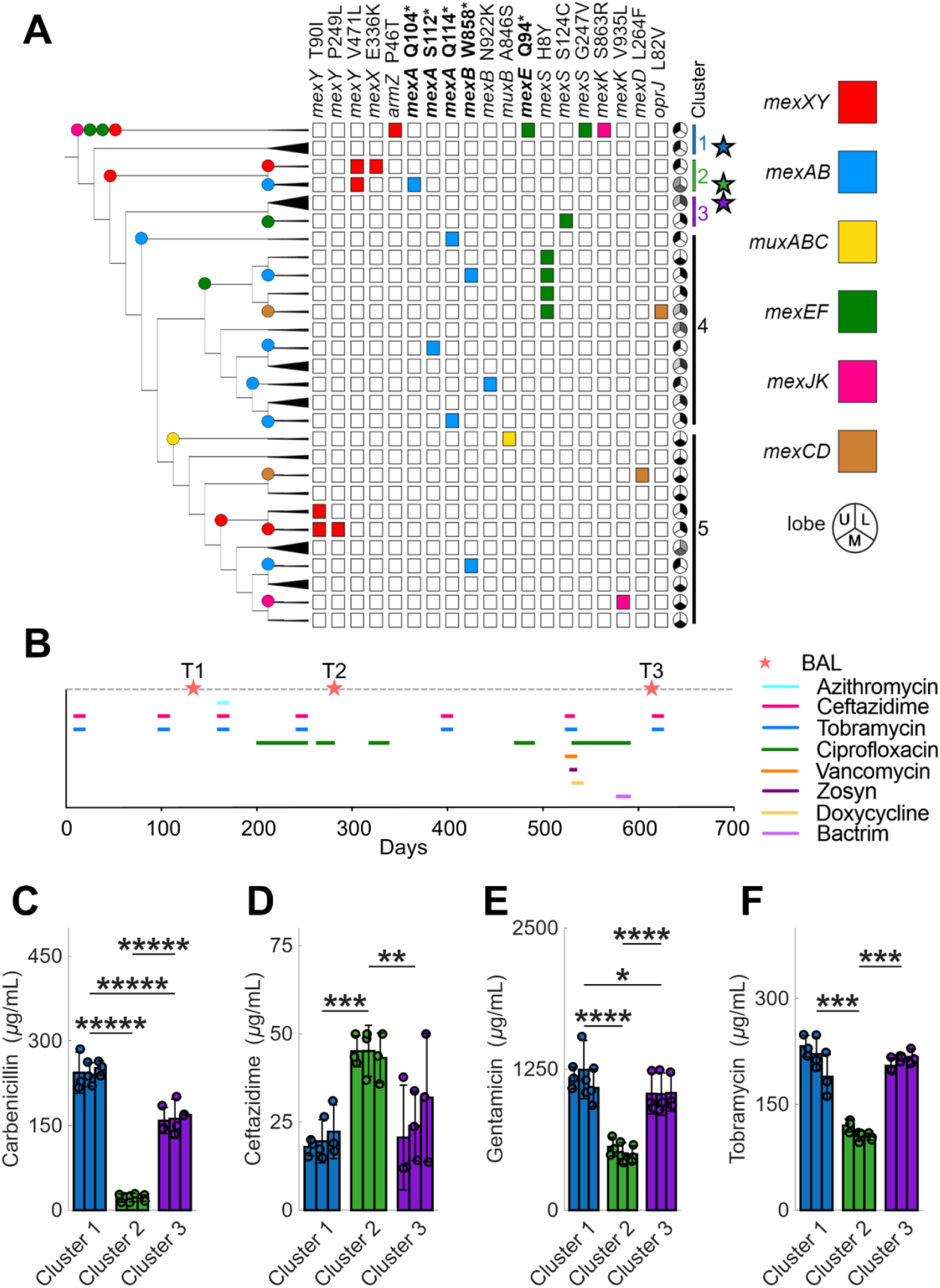
Efflux pumps are frequently mutated in CF isolates. **(A)** Mutations affecting efflux mechanisms are marked in the phylogeny tree of *P. aeruginosa* isolates. Adjacent strains with identical mutations were collapsed into the same leaf for clarity, with the size of the wedges representing the number of isolates collapsed into each leaf. Colored dots represent where each mutation likely occurred in the phylogeny tree. Bold x-axis labels represent nonsense mutations likely leading to loss of protein function. Cluster numbers are adjacent to the lobe pie graphs, and the stars denote which strains were chosen for the antibiotic resistance assays, shown in the subsequent panels. **(B)** Timeline of *P. aeruginosa* isolate collection and antibiotic usage over the study period. Peach stars denote the dates of bronchoalveolar lavage (BAL) sampling corresponding to timepoints T1, T2, and T3, and horizontal bars indicate periods of antibiotic administration. IC_50_ values were determined for isolates from three clusters for **(C)** carbenicillin, **(D)** ceftazidime, **(E)** gentamicin, and **(F)** tobramycin. These clusters are denoted by stars in Fig. 4A. For each cluster, three strains collected at Timepoint 1 from the right upper lobe were analyzed. No statistically significant differences were found between strains within the same clusters. Asterisks denote significance levels for pairwise comparison between clusters: ***** p < 10^-5^, **** p < 10^-4^, *** p < 10^-3^, ** p < 10^-2^, and * p < 0.05.

We observed many mutations in *mex* multidrug efflux pumps, *P. aeruginosa* main line of defense against antibiotics and other toxic compounds. We found three independent instances of mutations in *mexS,* a regulator of MexEF, which confers resistance to chloramphenicol, sulfamethoxazole, trimethoprim, and fluoroquinolones^39,40^. These mutations were present across the upper, middle, and lower lobes of the right lung, and some of the mutated residues were previously reported to increase antibiotic resistance^41^. We also found missense mutations in *mexXY* in two distantly related clusters, present in all lobes. MexXY is the primary efflux mechanism for aminoglycosides such as tobramycin, which is a first-line treatment for *P. aeruginosa* CF infections^42^. In both clusters, some lineages acquired sequential point mutations in the *mexXY* genes, suggesting adaptation of the efflux pump to specific drugs. Previous studies have shown that mutations in *mexXY* can enhance resistance to aminoglycosides^43^, but potentially at the cost of increased susceptibility to other antibiotics^44^. We also found 7 independent mutation events in *mexAB*, the primary efflux mechanism for β-lactams such as ceftazidime^45^, observed across all time points, lobes, and multiple clusters. Surprisingly, 6 of the 7 mutations lead to truncations, likely resulting in loss of protein function. Although MexAB and MexXY share partially overlapping substrate specificities, *mexAB* loss-of-function mutations are thought to increase MexXY activity by reducing competition for their shared outer membrane protein, OprM^46^. However, removing *mexB* did not affect aminoglycoside resistance in a past study^47^. Overall, these results indicate a strong selective pressure on efflux mechanisms affecting antibiotic resistance in the lung environment.

To determine if the evolved efflux pump alleles resulted in increased resistance to the antibiotics taken by the patient^45^, we measured resistance in select isolates to drug substrates of MexAB – carbenicillin and ceftazidime – and MexXY – gentamicin and tobramycin. We picked three isolates from each of neighboring Clusters 1, 2, and 3 **(Fig. 4A)**. All three isolates from Cluster 2 had a nonsense mutation in *mexA* and a missense mutation in *mexY*, while the isolates from Clusters 1 and 3 had no unfixed mutations in genes encoding multidrug efflux pumps. All nine isolates were collected from the right upper lobe at timepoint 1, following a course of ceftazidime and tobramycin treatment **(Fig. 4B)**. For each isolate, we determined both the IC_50_, which we define as the drug concentration required to half the growth rate of a log-phase liquid culture relative to the absence of drug (**Fig. 4C-F**), and the MIC, which we define as the drug concentration required for the final OD of the culture to remain under 0.1 after 20 hours (**Fig. S3**). For each analysis, we used fine drug gradients to allow the detection of small differences in resistance (**Fig. S4-S7**).

We observed significant differences in resistance between clusters, although strains within the same cluster showed no statistically significant differences in resistance to any of the antibiotics tested. For all four drugs, Cluster 2 strains differed significantly from Clusters 1 and 3, which were largely similar to each other. Despite the multiple mutations in *mexXY* and nonsense mutation in *mexA*, isolates from Cluster 2 displayed lower resistance to both aminoglycosides tested, compared to the other clusters. Strikingly, Cluster 2 isolates also exhibited nearly a tenfold decrease in carbenicillin resistance, likely due to the nonsense mutation in *mexA*. However, Cluster 2 isolates showed slightly higher resistance to ceftazidime, which was administered as a treatment and is also a substrate of MexAB, suggesting evolution of resistance through an alternative mechanism.

Together, these findings indicate that isolates from the same infection produce substantial variation in antibiotic resistance, even among isolates collected from the same lung lobe and timepoint. Of note, all isolates tested were more resistant to aminoglycosides than laboratory strains (**Fig. S6–S7**). However, it remains unclear how much of the antibiotic resistance can be explained by mutations in efflux pumps alone. While the mutation pattern in the isolates in Cluster 2 would suggest increased resistance to aminoglycosides and decreased resistance to β-lactams, the resistance profiles we measured for these strains do not support these expectations. Collectively, these observations suggest that additional mechanisms beyond *mex* efflux pumps play a major role in shaping the resistance profiles of the CF isolates.

## DISCUSSION

The goal of this study was to assess the genotypic diversity of a chronic infection population derived from a single strain type (**Fig. 1A**) in an individual with a mild-to-moderate CF lung disease within and across separate lobes of the right lung and over time. The infection population was caused by a strain belonging to MLST499 **(Fig. S1)**. The dimension of time for our sample set provides important clues as to how the population structure shifted over time.

Our phylogenetic analysis revealed groups of closely related isolates with good bootstrap support that were comprised entirely of isolates from a single lobe, supporting the compartmentalization hypothesis outlined in the study by Jorth, et al.^21^, but one cannot exclude the possibility that with the analysis of a larger number of strains, different lineages would be present in different regions of the lung. Our dataset also shows that individual lobes contain multiple lineages, and that specific lineages can be found across multiple lobes. In addition, while a particular genotype may dominate a particular lobe at a single time point, this same genotype may not persist in the same lobe or even in the overall population at subsequent time points (**Fig. 1C**).

Our analyses suggest that evolutionary rates were not uniform throughout the lung **(Fig. 2A**-Cluster 1). These differences may indicate that local factors, such as inflammation and the concomitant production of microbe damaging products or damage which may increase access to nutrients that may alter growth rate or population size, may impose distinct evolutionary pressures. In addition, specific unfixed mutations may impact mutation rate. We did not observe mutations in genes for which loss of function leads to a higher mutation rate.

We found that at singular time points, particular genotypes may indeed have been isolated to a single lobe of the lung **(Fig. 3B-C)**, and this finding in the context of mild-to-moderate disease resembles what has previously been described for more advanced lung disease settings^21,25^. However, when multiple time points were considered broad population shifts and changes in the location of particular genotypes hint that strict compartmentalization is not faithfully maintained over time **(Fig. 2A)**. Furthermore, migration rates were discontinuous in time with mixing occurring especially between the upper and the lower lobe around T2 followed by differentiation across lobes being rebuilt. It is important to note that there were changes in antibiotics with the introduction of ciprofloxacin also around T2 (**Fig. 4B**) suggesting that changes in disease state may have contributed to increased rates of migration. Because these populations were neither static nor isolated, they do not support a model in which differences in environment within specific lobes (e.g. drug access or extent of damage) was a major driver of evolution in this patient.

Our analyses found evidence of parallel mutations in different clusters and recent migration of genotypes across different lobes leading to their co-existence **(Fig. 3)**. One example was in the regulation of alginate production. While all isolates had identical loss-of-function mutations in *mucA,* which promotes mucoidy, we found multiple examples of mutations in *algU*, *algB*, and potentially other genes such as *algP* and *algL* that suppressed this phenotype **(Fig. 3A**, **Fig. 3E)**. The extensive genome sequence data generated in this study provide a valuable resource for future investigations into additional mutations that may influence the fitness of strains with altered activity within the Muc–Alg regulatory pathway. For instance, Cluster 1 strains with intact AlgB also carry different alleles of *migA*, which encodes a glycosyltransferase that modifies lipopolysaccharide (LPS), and PA4776, which encodes the sensor kinase PmrB, another regulator of LPS modification **(Supplementary Data 1)**^48,49^. Further, we found a Q45* *lasR* allele fixed in the population that is predicted to lead to loss-of-function. The ubiquity of this allele is interesting, as *lasR* loss-of-function mutations have been associated with worse patient outcomes^50,51^. Despite this, the individual with CF studied here has maintained relatively strong pulmonary function for someone chronically colonized by *P. aeruginosa*. Together, these findings highlight that the clinical impact of specific bacterial genotypes cannot be inferred from single mutations alone, and that disease severity likely reflects the combined effects of host factors, microbial community context, and the broader genetic background of the infecting population.

The results from our antibiotic resistance testing emphasize the need to culture many strains from an infection when determining an effective treatment protocol, as we demonstrate that different cell populations cultured from the same environment have significantly different antibiotic resistance profiles **(Fig. 4C-F)**. Apart from antibiotic resistance, mutations in efflux pumps affect other CF-relevant phenotypes, such as quorum sensing and virulence. For instance, loss of MexAB function, one of the most ubiquitous mutations in this study, can lead to overproduction and accumulation of quorum sensing molecules^46,52^, which has been suggested to increase tissue invasiveness in the CF lung^46^. Similarly, *mexEF* overexpression has been reported to reduce virulence due to attenuated quorum sensing^53^, and *mexXY* overexpression was shown to relieve oxidative stress. Indeed, mutations in multidrug resistance mechanisms often emerge even in patients that were never exposed to antibiotic treatments^54^, underscoring that these systems play broader physiological roles beyond drug efflux alone. Together, these observations indicate that remodeling of efflux mechanisms is a central component of *P. aeruginosa* adaptation to the lung environment. As a result, mutations affecting antibiotic resistance may arise in response to diverse selective pressures unrelated to clinical antibiotic use, complicating efforts to predict resistance phenotypes from genotype alone.

Chronic *P. aeruginosa* infections in the CF lung exist as dynamic and genetically diverse populations rather than uniform monocultures. As our findings demonstrate, even within a single species, the relative success of individual lineages can shift across both space and time. This heterogeneity has important implications in the context of antibiotic resistance and personalized medicine. A clearer understanding of the complexity of chronic infection populations can inform more effective treatment strategies, emphasizing the need for clinical approaches that account for the presence of diverse and evolving bacterial communities rather than relying on the characterization of a single representative isolate.

## MATERIALS AND METHODS

### Sample collection from human subject

The samples were collected through a study that was approved by the Institutional Review Board at Dartmouth-Hitchcock, protocol number 22781, and all methods were carried out in accordance with relevant guidelines and regulations. Following written informed consent, the person with CF then underwent flexible bronchoscopy. During the BAL procedures, the right lung was sampled in the order of upper lobe (RUL), middle lobe (RML), then lower lobe (RLL). Following the procedure, the person with CF was monitored until they were stable for discharge. An aliquot of the BAL fluid from the RUL was sent to the DHMC clinical microbiology laboratory for routine culture analysis and a separate aliquot was sent to our lab for isolate purification and banking as described below.

### Strains, media, and growth conditions

Strains were maintained in lysogeny broth (LB) medium, which consists of 10 g tryptone, 5 g NaCl, and 5 g Yeast Extract per liter distilled H_2_O or LB agar which also contains 15 g of Bacto-agar per liter. To examine colony morphologies, isolates were stamped from 96-well freezer stock plate onto Pseudomonas Isolation Agar (PIA) plates (Difco 292710) and incubated in a humidity chamber at 37°C for 48 hours. All isolates, in liquid culture and on solid plates, were grown at 37°C.

### Isolation of colonies from bronchoalveolar lavage and expectorated sputum

Immediately after sample collection, BAL fluid at different dilutions in phosphate buffered saline was spread directly onto PIA and blood agar plates with a metal spreading tool and incubated at 37°C for 48 hours. Individual colonies were picked from plates with well-isolated colonies, patched onto PIA, and grown for 36-48 hours. These colonies were then stamped into liquid LB in a 96-well plate via 48-pin replicator and allowed to grow for 48 hours. Subsequently, 100 µL of this culture was combined with 100 µL of 50% glycerol stock and frozen for storage at - 80°C.

### Specific PCR analysis and single gene sequencing

Epicentre Yeast DNA Extraction Kit was used for gDNA extraction, with omission of the RNase incubation step for select strains. For *lasR* and *mucA* amplification, Taq Phusion polymerase was used in a total reaction volume of 25 µl (17.5 µl H_2_O, 5 µl GC Buffer, 1.5 µl DMSO, 0.5 µl of DNTPs, forward primer, reverse primer, and Taq Phusion, plus 1 µl gDNA per tube). Primers were as follows: LasR_seq_1 5’ CAA ACG CTG CGG TCT ATT GTT AAG TG 3’, LasR_seq_2 (Jack Hammond), 5’ CAG TCG TTT CGA GAA TGG CGA GAA C 3’, mucA_forward_seq+amp_RVG, 5’ TTG AGT TAC GAA GAT ATC GCC ACC GTG 3’ and mucA_REVERSE_seq+amp_RVG, 5’ GCG CTC GTA GAC GAA GGT GCC 3’.The resulting PCR products were run on a 1% agarose gel with SYBR Green DNA visualization stain to ensure specific amplification. Subsequently, the PCR amplicon was purified with PCR Purification Kit by Qiagen and Sanger sequenced by the Dartmouth Molecular Biology Core Facility. The resulting sequences were aligned and analyzed with SnapGene against the PAO1 reference genome.

### Library preparation and sequencing

Strains were inoculated from the frozen plates into 96 well plates containing into 300 µL of LB and grown overnight at 37°C. Beckman GenFind V2 kits were used to extract genomic DNA, which was subsequently normalized using the Qubit dsDNA BR Assay kit. Nextera XT kits are used to construct libraries consisting of roughly 500 basepair DNA size selected fragments and were normalized with the Qubit dsDNA HS Assay kit. Bioanalyzer analysis was submitted to the Roy J. Carver Technology Center at the University of Illinois Urbana-Champaign. Libraries were sequenced with Illumina HiSeq2500 platform at the Roy J. Carver Technology Center to obtain 250 nucleotide paired end reads. An isolate was chosen for Nanopore sequencing at the same center to serve as a reference for intrapopulation mutation analyses.

### Reference genome assembly and multi-locus sequence type determination

Internal reference for isolate T1_RLL_B5 was assembled from Oxford Nanopore long reads and Illumina Hiseq2500 paired end reads for error correction under Unicycler was used under ‘--conservative’ mode for long read assembly. Annotation for internal reference genome was performed with Contigs of non-reference genomes were built from HiSeq reads with SPAdes assembler to explore gene content^55^. Settings for spades were performed under “—careful” settings with both forward and reverse fastq reads. Coverage of contigs ranged from 20X to 121X and N50 ranged from 2731-58765 bp within HiSeq2500 sequenced genomes. Annotation of Nanopore assemblies were performed with prokka^56^ parameters “--genus *Pseudomonas* --usegenus --species *aeruginosa* –rfam” and a PAO1 protein reference database.

Multi-locus sequence type (MLST) was identified by aligning reads to the PubMLST database using the software package ARIBA^57^. Multi-locus sequence types were identified using de novo contigs – assembled from Illumina short reads of T1_RLL_B5 using SPAdes software, --careful setting alignment – against the PAO1 reference genome using MUMmer software. T1_RLL_B5 sequences for the following MLST loci (“*acsA*” “*aroE*” “*guaA*” “*mutL*” “*nuoD*” “*ppsA*” “*trpE*”) were extracted and queried into PUBMLST.org to determine a the sequence type (ST). Loci sequences from the PAO1 reference were also submitted to validate workflow.

### Quality filtering and SNP analysis

After high-throughput sequencing of isolates, raw HiSeq reads were trimmed with trimmotatic with “2:36:10 LEADING:20 TRAILING:20 MAXINFO:50:0.97 MINLEN:50” settings^58^ and aligned to the internal reference. With samtools and breseq^31,59^, SNPs and indels that varied across the population were identified. After initial phylogenetic analyses of the SNP calls by breseq revealed a phylogenetic tree with low boot strap values and the appearance of rampant recombination, we undertook an unbiased filtering of our data to remove genomes and positions that were aberrant due to low or high read depth or low consensus scores. Consensus scores for SNPs were calculated as the sum of PHRED quality scores of the bases which support a SNP divided by the sum of all quality scores of bases mapped at that position. The filtering was performed as follows. Step 1: Isolates were removed if the average read depth was <30, as low read depth led to a high frequency of missed SNP calls (52 out of 527 isolate genomes removed). Step 2: For each genome, the number of positions with a consensus score of less than 0.9 was determined. Isolate genomes for which this value was more than 2 standard deviations higher than the average were removed (25 isolate genomes removed). Our analyses suggested samples with a high number of positions with a low consensus score likely contained contamination from another strain which impaired SNP calling even if present at a low level. Step 3: we removed positions that had an average read depth across isolates of >2 standard deviations from the mean across all genomes or a read depth less than ten (17 out of 721 positions removed). Step 4: For each position, the number of isolates with consensus score <0.9 was determined. Positions were removed if the number of isolates with a low consensus score at that position was more than two standard deviations away from the mean. (1 additional position removed). We carried forward 450 isolate genomes and 663 variable positions for further analyses. The impacts of SNPs were determined using SNPeff^60^. Code for consensus score determination is available at https://github.com/debhogan-208/Ritz_Clay_450_Pa_isolates.

### Main phylogenetic tree construction

For phylogenetic tree construction, nexus files were constructed using either MEGAX or Geneious Prime software after initial nucleotide alignment between sample core SNPs^61^. PAUP* tree building software^62^ was used to perform a heuristic parsimony search to find the best tree distance tree based on these core SNPs. Bootstrapping was performed with nreps=100 (“bootstrap nreps=100 search=heuristic/ addseq=random nreps=10 swap=tbr hold=1;”). iTOL was used to visualize trees and add metainformation^63^. To create trees from distant strains of *Pseudomonas*, a core genome was created with Spine against available assembled genomes using consensus alignments. MUMmer was used to align assemblies to a core genome sequence at default settings^64,65^. Delta-filter was applied to alignment to exclude SNPs found in repeats.

### Population level analysis

A series of R packages were used for population level analyses. Binary tables were used for mutations found between all samples within a subject containing lobe and date information. Data manipulation and import were performed with “dplyr” and “poppr” ^66^. Rarefaction curves were created with “vegan” package. Fst and Gst analysis were performed with StaMPP^67^.

### Phylogenetic analysis of *P. aeruginosa* migration patterns among lobes of the right lung

To examine the rate and heterogeneity of evolution among isolates, we performed a root-to-tip regression of genetic distance (**Fig. 1C**) against sample day^32^. The slope of this line approximates the average rate of evolution under a strict molecular clock model (**Fig. 2A**). To quantify the significance of this estimate, we used a permutation test in which the correlation between divergence and time was compared to a null distribution of the same statistic calculated with lobe locations of each tip shuffled for 10000 replicates^68^. The P value was calculated as the proportion of replicates in the null distribution with a correlation greater than or equal to the observed correlation.

Time-resolved phylogenetic trees were used to infer migration patterns of *P. aeruginosa* among the upper, middle, and lower lobes of the right lung using samples from BAL at days 128, 279, and 613. Nucleotides at 663 variable sites among isolates were concatenated to form input sequences. Markov Chain Monte Carlo was used to sample the posterior distribution of tree topologies, branch lengths, and node heights under a uncorrelated lognormally distributed relaxed molecular clock model implemented in BEAST 2.6.7^33,34^. Nucleotide substitution probabilities were modelled using the HKY substitution model^69^. A coalescent Bayesian skyline with 5 intervals was selected as the tree prior^70^. Migration among lobes was modelled using a 3 state asymmetric continuous time Markov “mugration” model implemented in the BEAST classic package v1.6.0^33,35^. MCMC was run for 1×10^9^ iterations, with a 10% burn-in. Convergence of all parameter estimates (excluding branch rate categories) was assessed as having effective sample size (ESS) values >200 using Tracer v1.7.3^71^. To reduce computational complexity, 9000 trees were re-sampled from this distribution, and a maximum clade credibility tree was used to visualize the posterior distribution of time trees with sampled lobe locations at each node (**Fig. 2B**). To represent migration events among lobes, switches among lobes, in which a child node had a different location than its parent node, were quantified for each sampled tree (**Fig. 2C**). Tree analyses were performed with custom scripts written R 4.5.1^72^ using the packages *ape* v5.8-1, *treeio* v1.34.0, *ggtree* v4.0.1, and *qgraph* v1.9.8^73–76^. Scripts and BEAST XML files to reproduce these analyses are available at https://github.com/hoehnlab/publication_scripts.

### Mucoid quantification

Each isolate was plated three times separately. Samples were diluted in PBS to yield between 30 and 150 colonies on PIA plates. The fraction of colonies with the mucoid morphology was assessed after 48 hours. Before statistical comparisons, the assumption of normal distributions was checked using the Shapiro-Wilk test. The normal distribution assumption was rejected for at least one within-clusters dataset for timepoint 3, but it was not rejected for the other timepoints. To account for this, the Wilcoxon Rank-Sum Test was used to compare the groups in timepoint 3 (**Fig. 3D**), and one-way ANOVA was performed using the Tukey-Kramer correction to complete pairwise comparisons for the other timepoints (**Fig. 3B-C**).

### Antibiotic resistance assays

All resistance assay experiments were conducted in a modified M63 minimal medium (10 g/L (NH_4_)_2_SO_4_, 68 g/L KH_2_PO_4_, 2.5 mg/L FeSO4·7H2O) supplemented with 2 g/L glucose, 1 g/L casamino acids, 0.12 g/L MgSO_4_, and 0.5 mg/L thiamine. Drug stock solutions were freshly made from powder stocks (Sigma Aldrich and Cayman Chemical Company). We assessed the IC_50_ and MIC of 3 different CF isolates from 3 separate clusters, as well as the lab strains PA14 and PAO1, for a total of 11 strains. Each strain had 3 biological replicates, and each replicate was assayed on a separate day. Freshly grown colonies on LB plates were inoculated into LB media and grown overnight, diluted 200-fold in 96-well microtiter plates (Falcon) containing 200 μL of fresh medium and antibiotic solution per well. After inoculation, OD (absorbance at 595 nm) was recorded by a plate reader (Synergy Neo-2) every 5 minutes for at least 20 hr. A concentration gradient of each antibiotic was set up over multiple 96-well plates. We used custom Matlab scripts to calculate the MIC, which we define as the drug concentration required to cause the final OD to be under 0.1 after 20 hours, as well as the IC_50_, which we define as the drug concentration that causes the resulting steady-state growth rate of the population to be halved relative to the no-drug condition. We used fine gradients and interpolated between data points to find both resistance measurements. The IC_50_ is calculated by linear regression of log_2_(OD) during the exponential growth phase (**Fig. S4-S7**). Any strains that did not cross the IC_50_ or MIC thresholds were assigned the highest drug concentration tested. Before statistical comparisons, the assumption of normal distributions was checked using the Shapiro-Wilk test. The normal distribution assumption was rejected for at least one within-clusters dataset, but it was not rejected for the between-clusters dataset. To account for this, Kruskal-Wallis was performed using the Bonferroni correction on strains within the same clusters, and one-way ANOVA was performed using the Tukey-Kramer correction on strains between clusters. When assessing statistical differences between the different clusters, antibiotic resistance measurements of biological replicates were first averaged together.

### Collapsed phylogenetic trees

The multidrug efflux pump and alginate collapsed phylogenetic trees were derived from the main phylogeny shown in **Fig. 1B**. Strains carrying unfixed non-synonymous mutations relative to the ancestor in the selected genes were identified and annotated with the strains’ lobe locations, cluster numbers, and the genes’ color categories. Adjacent strains sharing identical non-synonymous mutations were collapsed into the same leaf by pruning redundant branches. In the pruned trees, wedges at each leaf are proportional to the number of strains represented by that branch. In the multidrug efflux pump tree, stars denote which clusters were chosen to test in the antibiotic resistance assays. Transversion direction was determined by identifying the strand orientation of each gene of interest using online resources^77^. Code for construction of the trees is available at https://github.com/debhogan-208/Ritz_Clay_450_Pa_isolates.

### Grape tree

The grape tree was made from a minimum spanning network generated from the SNP alignment and visualized in GrapeTree version 1.5.^78^

## Acknowledgements

The authors thank Nora Grahl, Ph.D. and Alex Crocker, Ph.D. for assistance in collection of *P. aeruginosa* clinical isolates, and Jack Hammond, Ph.D. and Colleen Harty, Ph.D. for providing verified primers for Sanger sequencing of *lasR* and *algU*, respectively. We thank Laura Suttenfield, Ph.D. for assistance with data processing in R and J. Cristobal Vera, Ph.D. for early data analyses.

This research was supported by the Cystic Fibrosis Foundation Research Development Program (BOMBER24G0) to D.A.H. and A.A. through the DartCF Respiratory Research Core. Other support for D.S. and D.R. came from NSF grants PHY-2412766, DMS-2527337 and U.S. Department of Energy grant DE-SC0026232. A.A. was further supported by NIH grant R01HL174700. K.B.H. was supported by National Institute for General Medical Sciences grant R35GM160119. James O. Freeman Presidential Scholar Fund provided support to R.V.G. This research was supported by funding from the CFF foundation and the Paul G. Allen Family Foundation (R.J.W.) and the Thomas S. Kosasa endowment (D.A.H.). Equipment and tools from BioMT were used, which is supported through NIH NIGMS grant P20-GM113132.

---

**Supplemental Dataset S1.** Mutations identified among the 450 isolates sequenced from the subject. SNPs were called relative to the internal reference sequence which was derived from isolate T1_RLL_B5.

**Supplemental Dataset S2.** Indels identified among the 450 isolates sequenced from the subject. Polymorphisms were called relative to the internal reference sequence which was derived from isolate T1_RLL_B5.

